# The plant pathogen *Pectobacterium atrosepticum* contains a functional formate hydrogenlyase-2 complex

**DOI:** 10.1101/688135

**Authors:** Alexander J. Finney, Rebecca Lowden, Michal Fleszar, Marta Albareda, Sarah J. Coulthurst, Frank Sargent

**Author notes:** **Corresponding author:** Prof Frank Sargent FRSE, School of Natural & Environmental Sciences, Newcastle University, Devonshire Building, NEWCASTLE UPON TYNE NE1 7RU, England.

## Abstract

*Pectobacterium atrosepticum* SCRI1043 is a phytopathogenic gram-negative enterobacterium. Genomic analysis has identified that genes required for both respiration and fermentation are expressed under anaerobic conditions. One set of anaerobically expressed genes is predicted to encode an important but poorly-understood membrane-bound enzyme termed formate hydrogenlyase-2 (FHL-2), which has fascinating evolutionary links to the mitochondrial NADH dehydrogenase (Complex I). In this work, molecular genetic and biochemical approaches were taken to establish that FHL-2 is fully functional in *P. atrosepticum* and is the major source of molecular hydrogen gas generated by this bacterium. The FHL-2 complex was shown to comprise a rare example of an active [NiFe]-hydrogenase-4 (Hyd-4) isoenzyme, itself linked to an unusual selenium-free formate dehydrogenase in the final complex. In addition, further genetic dissection of the genes encoding the predicted membrane domain of FHL-2 established surprisingly that the majority of genes encoding this domain are not required for physiological hydrogen production activity. Overall, this study presents *P. atrosepticum* as a new model bacterial system for understanding anaerobic formate and hydrogen metabolism in general, and FHL-2 function and structure in particular.

**Significance Statement:** *Pectobacterium atrospecticum* contains the genes for the formate hydrogenlyase-2 enzyme, considered the ancient progenitor of mitochondrial respiratory Complex I. In this study, the harnessing of *P. atrosepticum* as a new model system for understanding bacterial hydrogen metabolism has accelerated new knowledge in FHL-2 and its component parts. Importantly, those component parts include an unusual selenium-free formate dehydrogenase and a complicated [NiFe]-hydrogenase-4 with a large membrane domain. FHL-2 is established as the major source of molecular hydrogen produced under anaerobic conditions by *P. atrospectium*, however surprisingly some components of the membrane domain were not essential for this activity.

## Introduction

Many members of the *γ*-Proteobacteria are facultative anaerobes with the ability to switch their metabolisms to exploit the prevailing environmental conditions. Aerobic or anaerobic respiration is generally preferred, depending on the availability of respiratory electron acceptors. In this phylum, and specifically under anaerobic conditions, the three-carbon product of glycolysis, pyruvate, is often further metabolised by the oxygen-sensitive pyruvate formatelyase enzyme to generate acetyl CoA and the one-carbon compound formic acid (Pinske & Sawers, 2016). Studies of the model bacterium *Escherichia coli* have established that endogenously-produced formate is initially excreted directly from the cell using a dedicated channel (Lu *et al.*, 2011). Under respiratory conditions this formate would be used as an electron donor through the activity of periplasmic enzymes, but under fermentative conditions the formate accumulates in the extracellular milieu until its rising concentration begins to cause a drop in extracellular pH. This is thought to trigger formate re-uptake, which in turn induces synthesis of formate hydrogenlyase (FHL) activity in the cell (Rossmann *et al.*, 1991, McDowall *et al.*, 2014, Sargent, 2016). FHL activity then proceeds to detoxify the formic acid by disproportionation to carbon dioxide and molecular hydrogen (H_2_).

While FHL activity has been characterised in *E. coli* (Sargent, 2016), it is not confined to enteric bacteria and has been reported across the prokaryotic domains, including in hyperthermophilic archaea where it is not only involved in pH homeostasis but also in generating transmembrane ion gradients (Kim *et al.*, 2010, Lim *et al.*, 2014). The ion pumping activity stems from an evolutionary link between FHL and the respiratory NADH dehydrogenase Complex I (Bohm *et al.*, 1990, Batista *et al.*, 2013, Schut *et al.*, 2016). Like Complex I, FHL comprises a cytoplasmic catalytic domain linked to an integral membrane domain. The catalytic domain contains a [NiFe]-hydrogenase of the ‘Group 4’ type, which is primarily dedicated to H_2_ production (Greening *et al.*, 2015), and is linked by [Fe-S]-cluster-containing proteins to a molybdenum-dependent formate dehydrogenase (Maia *et al.*, 2015). The FHL membrane domain is predicted to take two different forms allowing the enzyme to be further sub-classified as either ‘FHL-1’ or ‘FHL-2’ (Sargent, 2016, Finney & Sargent, 2019). The FHL-1 is the predominant archetypal FHL activity of *E. coli* (McDowall *et al.*, 2014) and comprises [NiFe]-hydrogenase-3 (Hyd-3), formate dehydrogenase-H (FdhF), and a relatively small membrane domain compared to Complex I that contains only two proteins (Figure 1A). Genes for the much less well understood FHL-2 enzyme are also found in *E. coli* (Andrews *et al.*, 1997). This isoenzyme is predicted to comprise a [NiFe]-hydrogenase-4 (Hyd-4), an as-yet undefined formate dehydrogenase, and a much larger membrane domain than FHL-1, containing at least five individual integral membrane subunits and more closely resembling the Complex I structure (Figure 1B). Understanding the structure, function and physiological role of *E. coli* Hyd-4 and FHL-2 has been hindered by poor native expression levels (Skibinski *et al.*, 2002, Self *et al.*, 2004); a missing important accessory gene from the *hyf* cluster (Sargent, 2016); and a lack of consensus on the appropriate experimental conditions to test (Bagramyan *et al.*, 2001, Mnatsakanyan *et al.*, 2004).

**Figure 1:**
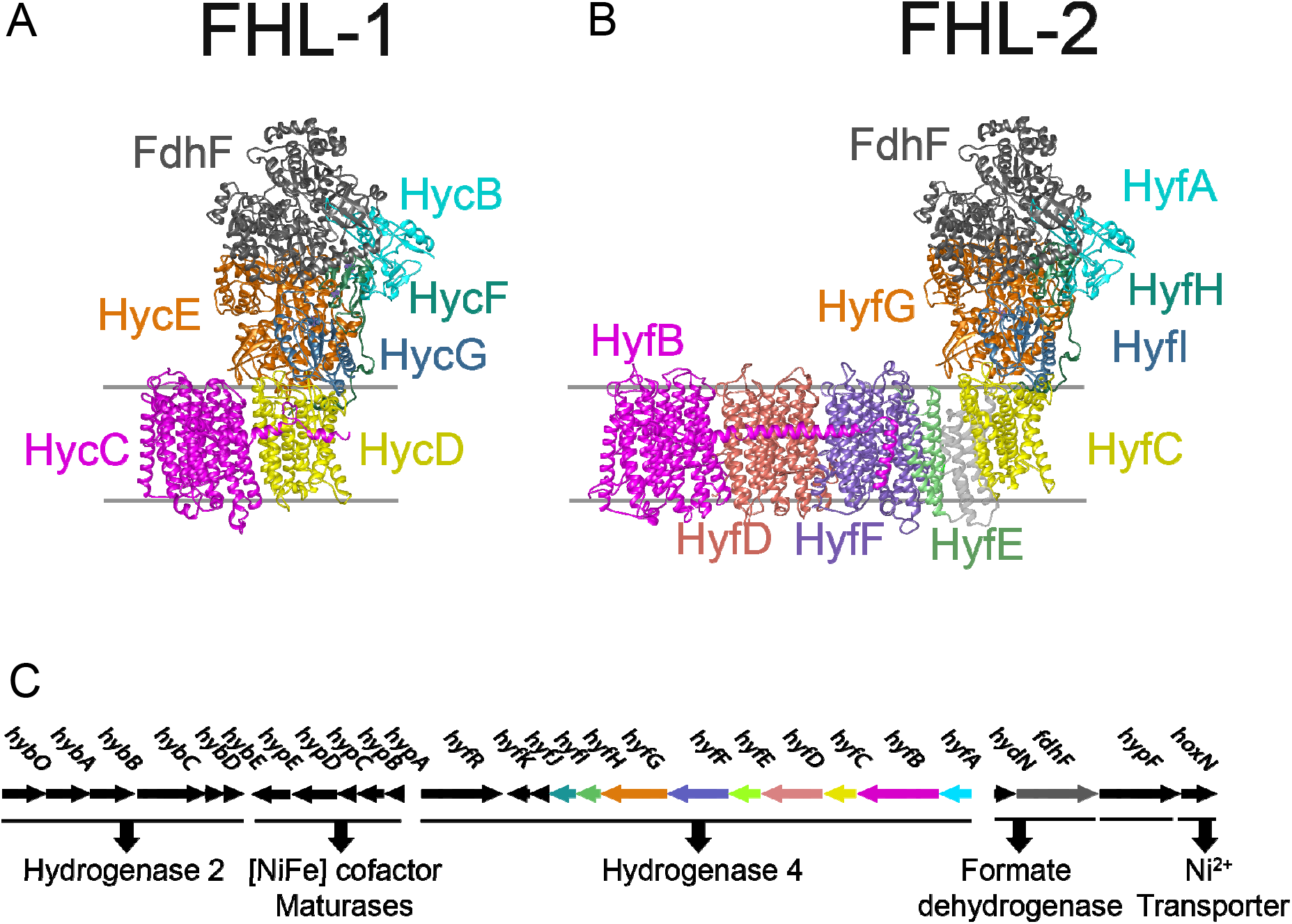
Biochemistry and genetics of formate hydrogenlyase. Structural models of **(A)** formate hydrogenlyase-1 (FHL-1) from *Escherichia coli* and **(B)** formate hydrogenlyase-2 (FHL-2) from *Pectobacterium atrosepticum*. Subunits related at the primary and tertiary levels are coloured similarly. Figures were generated using Chimera (Pettersen *et al.*, 2004). **(C)** The genetic organisation of the hydrogen metabolism gene cluster of *P. atrosepticum* (ECA1225-ECA1252). Predicted gene product functions are indicated and the operon for Hyd-4 is colour coded to match the structure model in panel (B).

In order to bring fresh impetus to understanding and harnessing the FHL-2 complex, it was considered that an appropriate alternative biological model system was required. *Pectobacterium atrosepticum* SCRI1043 is a phytopathogenic *γ*-Proteobacterium that can grow under anaerobic conditions. A transcriptomic study identified a chromosomal locus (Figure 1C) that was transcribed under anaerobic conditions in this organism (Babujee *et al.*, 2012, Bell *et al.*, 2004). This locus neatly collects together almost all of the known genes for hydrogen metabolism (Figure 1C), including genes for a bidirectional Hyd-2-type [NiFe]-hydrogenase; genes for specialist metallo-cofactor biosynthesis; a putative formate-responsive transcriptional regulator; a predicted formate dehydrogenase gene; and an 11-cistron operon apparently encoding a Hyd-4 isoenzyme and its associated accessory proteins (Babujee *et al.*, 2012).

In this work, a molecular genetic approach was taken to characterise the hydrogen metabolism locus of *P. atrosepticum*. A bank of un-marked and in-frame gene deletion mutants was constructed and used to demonstrate unequivocally that the unusual FHL-2 is functional in *P. atrosepticum* and responsible for the majority of H_2_ production under anaerobic conditions. The complex was shown to contain an active Hyd-4 and, unusually, a version of formate dehydrogenase that does not rely on selenocysteine. Surprisingly, it was shown that many of the genes encoding the large membrane domain of FHL-2 can be removed without adversely affecting H_2_ production activity. Overall, this work introduces *P. atrosepticum* as a tractable model system and presents important genetic, biochemical and physiological characterisation of FHL-2 and [NiFe]-hydrogenase-4.

## Results

### P. atrosepticum *produces molecular hydrogen under anaerobic conditions*

*P. atrosepticum* SCRI1043 (Bell *et al.*, 2004) contains the genes for potentially H_2_-evolving enzymes (Babujee *et al.*, 2012). Therefore, the initial goal of this study was to establish the growth conditions under which molecular hydrogen could be evolved. First, the SCRI1043 wild-type strain was grown under anaerobic fermentative conditions in a minimal medium supplemented with 0.8% (w/v) glucose. The culture headspace was sampled at periodic intervals and the amount of H_2_ present quantified by gas chromatography (GC). Under these conditions, H_2_ evolution activity was found to be temperature dependent, with H_2_ accumulation in the headspace observed to be maximal when the phytopathogen was incubated at 20 or 24 °C (Figure 2A). Taking forward 24 °C as standard incubation temperature, H_2_ evolution was observed and found to level off after 40 hours incubation (Figure 2B).

**Figure 2:**
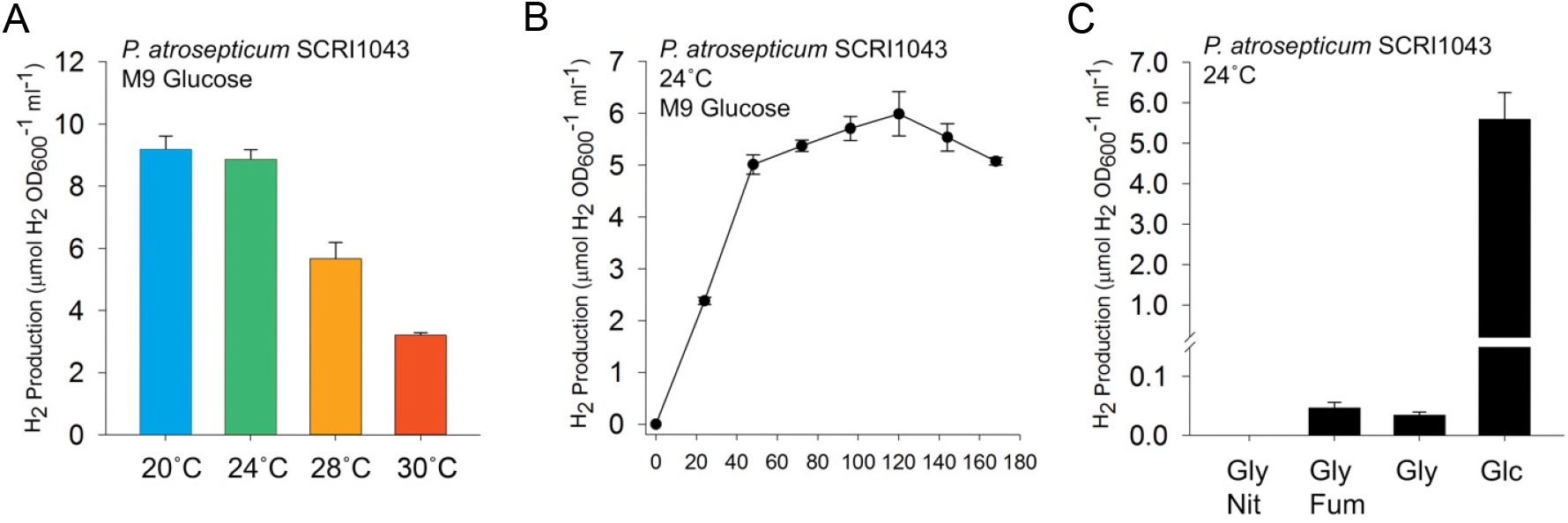
*P. atrosepticum* produces molecular hydrogen gas. **(A)** Anaerobic hydrogen production is optimal at lower temperatures. The *P. atroscpeticum* SCRI1043 parent strain was incubated in M9 medium supplemented with 0.8% (w/v) glucose for 168 hours at the temperatures indicated before gaseous H_2_ accumulation was quantified. **(B)** A time-course of H_2_ accumulation. *P. atrosepticum* SCRI1043 was incubated in M9 medium supplemented with 0.8% (w/v) glucose at 24 °C and gaseous H_2_ accumulation was measured every 24 hours. **(C)** *P. atrosepticum* SCRI1043 was incubated in M9 medium supplemented with either 0.5% (v/v) glycerol and 0.4% (w/v) nitrate (‘Gly Nit’); 0.5% (v/v) glycerol and 0.4% (w/v) fumarate (‘Gly Fum’); 0.5% (v/v) glycerol only (Gly); or 0.8% (w/v) glucose only (‘Glc’) at 24 °C for 48 hours. In all cases, the levels of molecular H_2_ in the culture headspace were quantified by GC and normalised to OD_600_ and culture volume. Error bars represent SD (*n* = 3).

When anaerobic respiratory conditions were tested, comprising 0.5% (v/v) glycerol and 0.4% (w/v) nitrate, H_2_ production was found to cease with no H_2_ detectable after 48 hours growth (Figure 2C). However, replacement of nitrate with 0.4% (w/v) fumarate as a terminal electron acceptor allowed the generation of low, but detectable, levels of H_2_ (Figure 2C). Maximal H_2_ production is observable under fermentative conditions (Figure 2C).

### Hyd-4 is the predominant hydrogen producing enzyme in P. atrosepticum

To determine the molecular basis of the observed H_2_ production activity, a molecular genetic approach was taken. Initially, the genes encoding the catalytic subunits of the [NiFe]-hydrogenases were targeted. First, a strain PH001 (Table 1) was constructed carrying an unmarked in-frame deletion of the *hyfG* gene, predicted to encode the catalytic subunit of a Hyd-4 isoenzyme. When cultured fermentatively in the presence of glucose, the PH001 (Δ*hyfG*) strain produced less than 5% of the total H_2_ accumulated by the wild-type control under the same conditions (Figure 3A). Next, the gene encoding the catalytic subunit of Hyd-2 (*hybC*) was tested. Mutant strain PH002 (Table 1) was prepared carrying only a Δ*hybC* allele and, in this case, H_2_ evolution under fermentative conditions was essentially indistinguishable from the wild-type strain (Figure 3A). Finally, a Δ*hybC* Δ*hyfG* double mutant (PH003, Table 1) was constructed and was found to be completely devoid of the ability to produce gaseous H_2_ (Figure 3A).

**Table 1:** *P. atrosepticum* strains and plasmids used in this study.

| Strain | Relevant genotype | Genomic Identifier | Source |
| --- | --- | --- | --- |
| SCRI1043 | - |  | Bell *et al.* 2004 |
| PH001 | $\Delta hyfG$ | ECA1241 | This work |
| PH002 | $\Delta hybC$ | ECA1228 | This work |
| PH003 | $\Delta hybC \Delta hyfG$ | ECA1228, ECA1241 | This work |
| PH004 | $\Delta fdhF$ | ECA1250 | This work |
| PH005 | $\Delta hybC \Delta fdhF$ | ECA1228, ECA1250 | This work |
| PH007 | $\Delta hybC \Delta hyfB-F$ | ECA1228, ECA1246-2 | This work |
| PH008 | $\Delta hybC \Delta hyfD-F$ | ECA1228, ECA1244-2 | This work |
| PH009 | $\Delta hybC hyfG^{His}$ | ECA1228, ECA1241 | This work |
| PH010 | $\Delta hypF$ | ECA1251 | This work |
| PH011 | $\Delta hoxN$ | ECA1252 | This work |
| PH013 | $\Delta hybC \Delta fdhD$ | ECA1228, ECA0093 | This work |
| PH015 | $\Delta hybC \Delta hyfR$ | ECA1228, ECA1236 | This work |
| PH018 | $\Delta hybC \Delta ECA1964$ | ECA1228, ECA1964 | This work |
| PH019 | $\Delta hybC \Delta fdhF \Delta ECA1964$ | ECA1228, ECA1250, ECA1964 | This work |
| PH020 | $\Delta hybC hyfG^{His} \Delta hyfB-F$ | ECA1228, ECA1241, ECA1246-2 | This work |
| PH021 | $\Delta hybC hyfG^{His} \Delta hyfD-F$ | ECA1228, ECA1241, ECA1244-2 | This work |
| PH027 | $\Delta hybC \Delta ECA1507$ | ECA1228, ECA1507 | This work |
| PH028 | $\Delta hybC \Delta fdhF \Delta ECA1507$ | ECA1228, ECA1250, ECA1507 | This work |

| Plasmid | Description | Source |
| --- | --- | --- |
| pSUPROM | Vector for expression of genes under control of the *E. coli* *tatA* promoter (Kan<sup>R</sup>) | Jack *et al.* 2004 |
| pSUPROM-*hyfG* | As pSUPROM with *P. atrosepticum hyfG* (ECA1241) | This work |
| pSUPROM-*hybC* | As pSUPROM with *P. atrosepticum hybC* (ECA1228) | This work |
| pSUPROM-*fdhF* | As pSUPROM with *P. atrosepticum fdhF* (ECA1250) | This work |
| pSUPROM-ECA1507 | As pSUPROM with *P. atrosepticum ECA1507* | This work |
| pSUPROM-ECA1964 | As pSUPROM with *P. atrosepticum ECA1964* | This work |
| pSUPROM-*fdhD* | As pSUPROM with *P. atrosepticum fdhD* (ECA0093) | This work |
| pSUPROM-*hypF* | As pSUPROM with *P. atrosepticum hypF* (ECA1251) | This work |
| pSUPROM-*hoxN* | As pSUPROM with *P. atrosepticum hoxN* (ECA1252) | This work |
| pSUPROM-*hyfR* | As pSUPROM with *P. atrosepticum hyfR* (ECA1236) | This work |

**Figure 3:**
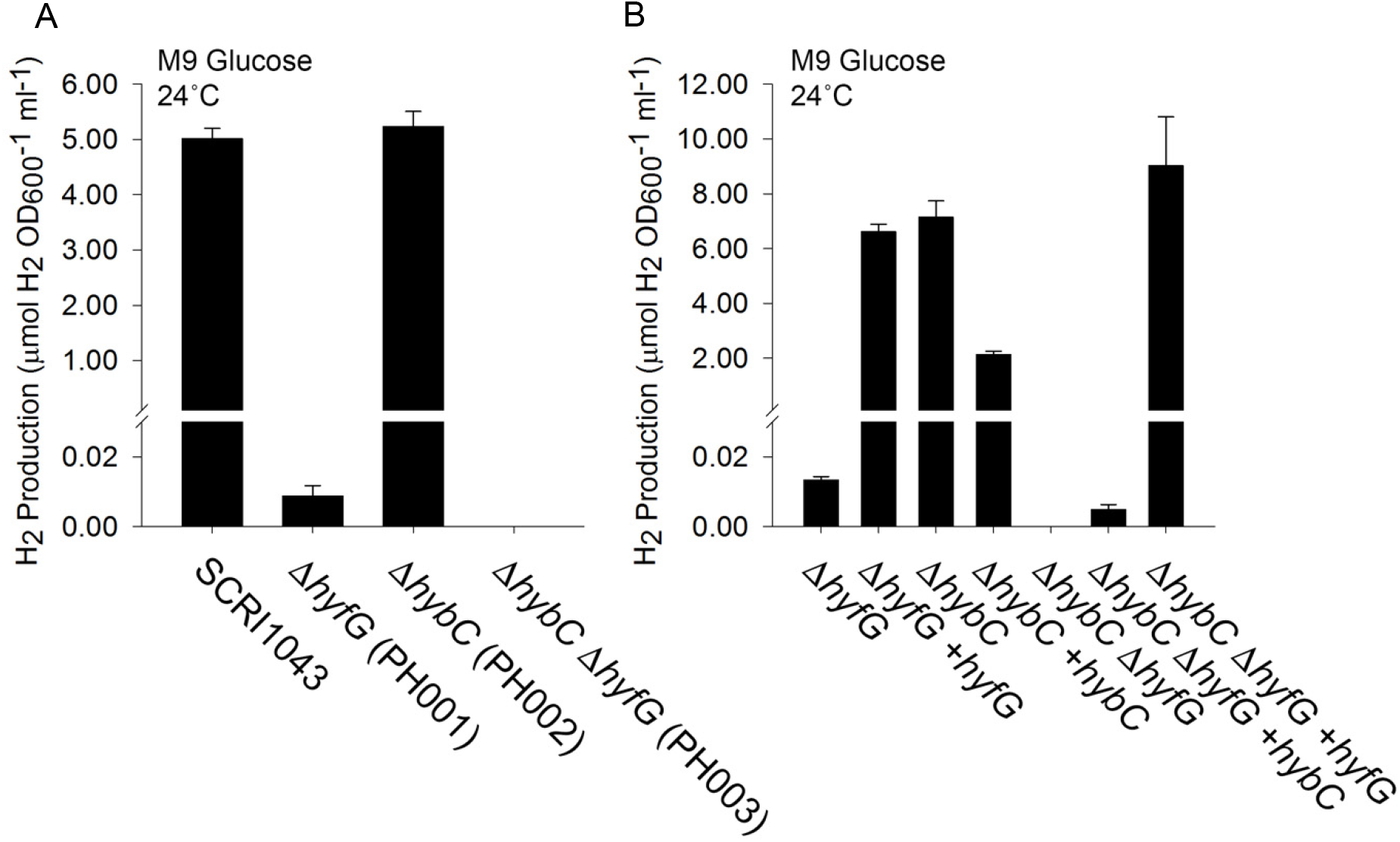
Hydrogen gas is produced by the activity of [NiFe]-Hydrogenase-4. **(A)** Hyd-4 is responsible for fermentative H_2_ production. *P. atrosepticum* parental strain SCRI1043 and mutants PH001 (Δ*hyfG*), PH002 (Δ*hybC*) and PH003 (Δ*hybC* Δ*hyfG*) were incubated in M9 medium supplemented with 0.8% (w/v) glucose at 24 °C for 48 hours. **(B)** Complementation of the mutant phenotype *in trans*. Strains PH001 (Δ*hyfG*), PH002 (Δ*hybC*) and PH003 (Δ*hybC* Δ*hyfG*) were separately transformed with plasmids encoding either HyfG or HybC under the control of constitutive promoters. Levels of molecular H_2_ in the culture headspace were quantified by GC and normalised to OD_600_ and culture volume. Error bars represent SD (*n* = 3).

*P. atrospeticum* can be stably transformed and plasmids encoding either *hyfG* or *hybC* were constructed. In the case of PH001 (Δ*hyfG*) and PH003 (Δ*hybC* Δ*hyfG*), H_2_ evolution could be rescued in the mutant strains by supplying extra copies of *hyfG* on a plasmid (Figure 3B).

Taken altogether, the data presented in Figures 2 **and** 3 demonstrate that Hyd-4 is responsible for the majority of physiological H_2_ production by *P. atrospeticum*, and that this activity is present under fermentative conditions at temperate growth temperatures ≤24 °C.

### P. atrosepticum contains an active FHL-2 with a selenium-free formate dehydrogenase

Having established that Hyd-4 was active, the next task was to test the hypothesis that Hyd-4 could be part of a wider FHL-2 complex (Figure 1). First, formate-dependence on H_2_ production was tested by growing the wild-type parental strain, the PH001 (Δ*hyfG*) strain, and the PH002 (Δ*hybC*) strain anaerobically in the presence of increasing amounts of exogenous formate (Figure 4A). A correlation was observed between the amount of H_2_ produced and the amount of formate added to the growth medium (Figure 4A), and H_2_ production remained dependent upon the presence of an active Hyd-4 (Figure 4A), providing initial evidence for a link between formate and H_2_ metabolism.

**Figure 4:**
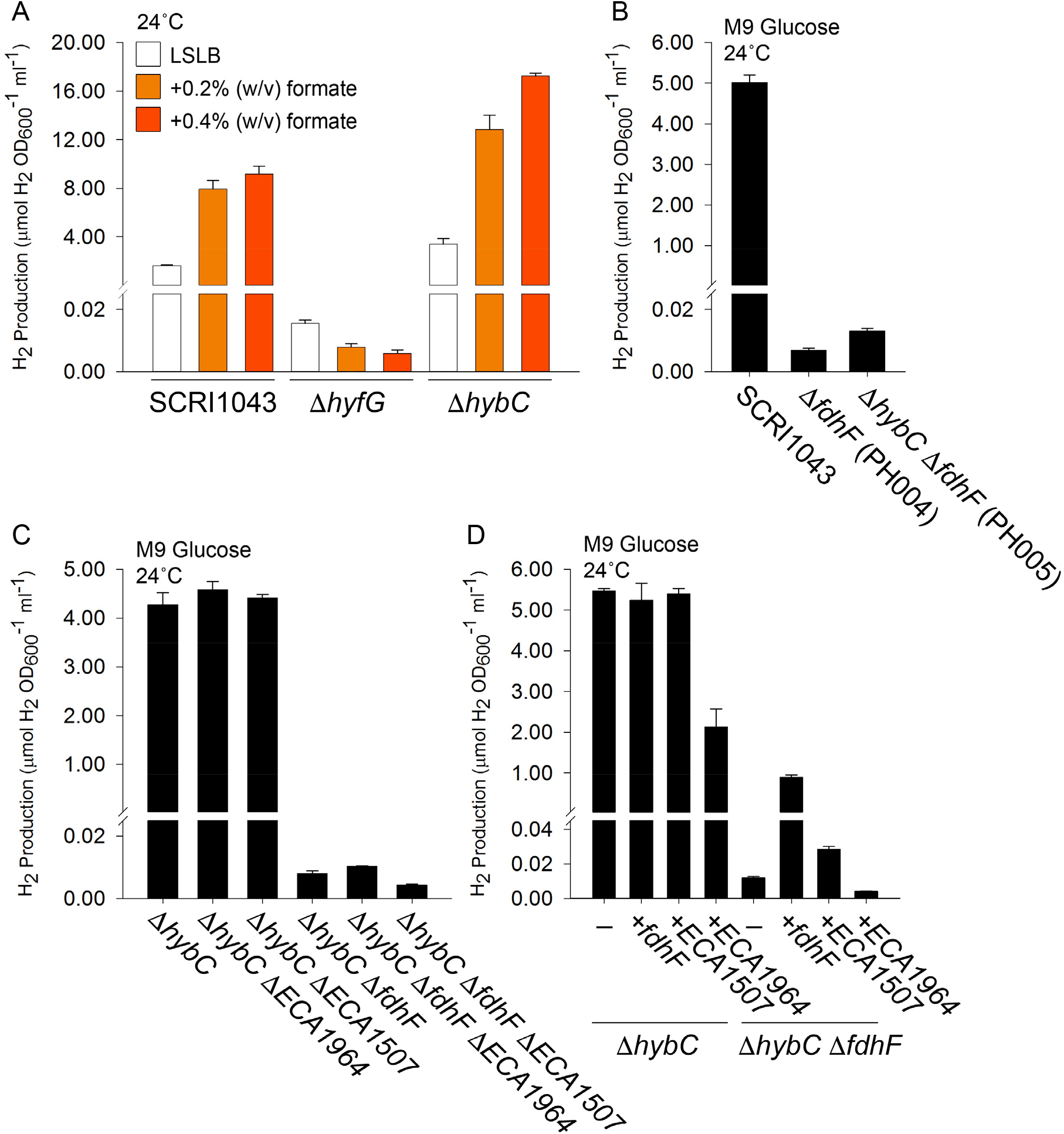
Hydrogen gas is produced by the activity of a selenium-free formate dehydrogenase. **(A)** Addition of exogenous formate increases H_2_ production. *P. atrosepticum* parental strain SCRI1043 and mutants PH001 (Δ*hyfG*) and PH002 (Δ*hybC*) were incubated in low-salt (5g/l) LB (LSLB) rich medium supplemented with 0.2% or 0.4% (w/v) formate at 24 °C for 48 hours. **(B)** The formate dehydrogenase encoded within the gene cluster is responsible for FHL-2 activity. Strains SCRI1043, PH004 (Δ*fdhF*), PH005 (Δ*hybC* Δ*fdhF*) were incubated in M9 medium supplemented with 0.8% (w/v) glucose at 24 °C for 48 hours. **(C)** Alternative formate dehydrogenase homologues do not have a major role in H_2_ production. Strains SCRI1043, PH002 (Δ*hybC*), PH019 (Δ*hybC* Δ*ECA1964*), PH028 (Δ*hybC* Δ*ECA1507*) and PH005 (Δ*hybC* Δ*fdhF*) were incubated in M9 medium supplemented with 0.8% (w/v) glucose at 24 °C for 48 hours. **(D)** Complementation of the mutant phenotype *in trans*. Strains PH002 (Δ*hybC*) and PH005 (Δ*hybC* Δ*fdhF*) were separately transformed with plasmids encoding either FdhF, ECA1964 or ECA1507 under the control of constitutive promoters. In all cases, the levels of molecular H_2_ in the culture headspace were quantified by GC and normalised to OD_600_ and culture volume. Error bars represent SD (*n* = 3).

The *P. atrosepticum* SCRI1043 genome contains a gene encoding a putative formate dehydrogenase close to those for Hyd-4 (Figure 1C). The gene product shares 85% overall sequence identity with *E. coli* FdhF but interestingly contains a cysteine residue at position 140 (**Supp. Figure S1**), which is occupied by a critical selenocysteine in the *E. coli* version and other related enzymes (Axley *et al.*, 1991). A mutant strain was therefore constructed (PH004, Table 1) carrying an Δ*fdhF* allele. The PH004 (Δ*fdhF*) strain produced very low levels of H_2_ gas under fermentative conditions (Figure 4B). Addition of a Δ*hybC* allele to the Δ*fdhF* strain to generate a double mutant (PH005, Table 1) had no further effect on the amount of H_2_ that could be produced (Figure 4B).

Elsewhere on the *P. atrosepticum* SCRI1043 genome two further homologs of FdhF are encoded. The *ECA1507* gene encodes a protein with 65% overall sequence identity with FdhF, and the *ECA1964* gene encodes a protein with 22% overall sequence identity with FdhF (Bell *et al.*, 2004). Deletion of the genes encoding *ECA1507* or *ECA1964* alone (Table 1) had no influence on the H_2_ production capability of the bacterium (Figure 4C). Moreover, when the genes were supplied in multicopy on an expression vector, neither was able to rescue the phenotype of the Δ*fdhF* mutant (Figure 4D).

Taken altogether, these data establish that *P. atrosepticum* SCRI1043 has functional formate hydrogenlyase activity where molecular hydrogen production is clearly linked to both formate availability and the presence of a formate dehydrogenase gene. Importantly, the predominant electron donor for the reaction is an unusual version of formate dehydrogenase that does not require selenocysteine at its active site, and the enzyme responsible for proton reduction is a [NiFe]-hydrogenase-4.

### The role of the FHL-2 membrane domain in hydrogen production

One clear defining structural difference between the FHL-1 type formate hydrogenlyase found in *E. coli* and the FHL-2 type of *P. atroscepticum* SCRI1043 is the number of genes encoding components of the membrane domains (Figure 1). An FHL-1 enzyme is predicted to contain two different membrane proteins, HycC (related to HyfB in FHL-2) and HycD (related to HyfC) (Figure 1A). Alternatively, an FHL-2 enzyme is predicted to contain three additional membrane proteins, including HyfE (not present in FHL-1) and two further homologs of HycC/HyfB, namely HyfD and HyfF (Figure 1B).

To explore the roles of the extra *hyfDEF* genes located within the FHL-2 locus, mutant strains were constructed (Table 1). First, versions of the Δ*hybC* strain PH002, lacking either the genes encoding the entire FHL-2 membrane domain (PH007: Δ*hybC*, Δ*hyfBCDEF* – Table 1) or lacking the extra membrane components not found in FHL-1 (PH008: Δ*hybC*, Δ*hyfDEF* – Table 1) were constructed. In addition, the Δ*hybC* strain PH002, producing Hyd-4 as the only active hydrogenase, was modified by addition of a 10-His sequence between codons 82 and 83 of the *hyfG* gene. This new epitope-tagged strain was called PH009 (Δ*hybC*, *hyfG*^His^ – Table 1). Finally, versions of PH009 lacking either the genes encoding the entire FHL-2 membrane domain (PH020: Δ*hybC*, *hyfG*^His^, Δ*hyfBCDEF* – Table 1) or lacking the extra membrane components not found in FHL-1 (PH021: Δ*hybC*, *hyfG*^His^, Δ*hyfDEF* – Table 1) were constructed.

Deletion of the genes encoding the entire membrane domain reduced the FHL-2-dependent H_2_ accumulation levels to around 5% of that observed in the parent strain (Figure 5A). The addition of the 10-His tag to HyfG allowed the Hyd-4 catalytic subunit to be visualised in whole cell extracts by Western immunoblotting (Figure 5B). The polypeptide was clearly detectable when *P. atroscepticum* was cultured under anaerobic fermentative conditions (Figure 5C). Interestingly, the amount of cellular HyfG^His^ was seen to increase when the genes encoding the membrane domain were removed (Figure 5). This is particularly pertinent for the PH020 strain (Δ*hybC*, Δ*hyfBCDEF*), which is essentially devoid of FHL-2 activity (Figure 5A), since it can be concluded that genetic removal of the complete membrane domain does not destabilise the Hyd-4 catalytic subunit, but instead leads to a physiologically inactive enzyme. It is also notable that in the absence of the genes encoding membrane proteins that the HyfG^His^ protein migrates as two electrophoretic species during SDS-PAGE (Figure 5B). This is a common observation for catalytic subunits of [NiFe]-hydrogenases as they are synthesised as precursors that undergo proteolytic processing at the C-terminus following cofactor insertion (Bock *et al.*, 2006). In this case, the faster migrating species was calculated as 56.4 kDa, while the slower migrating species was estimated as 62.6 kDa by SDS-PAGE. The predicted molecular mass of HyfG^His^ prior to proteolytic processing is 67,559 Da, and the predicted molecular weight of the 32-residue C-terminal tail that is removed is 3,821 Da.

**Figure 5:**
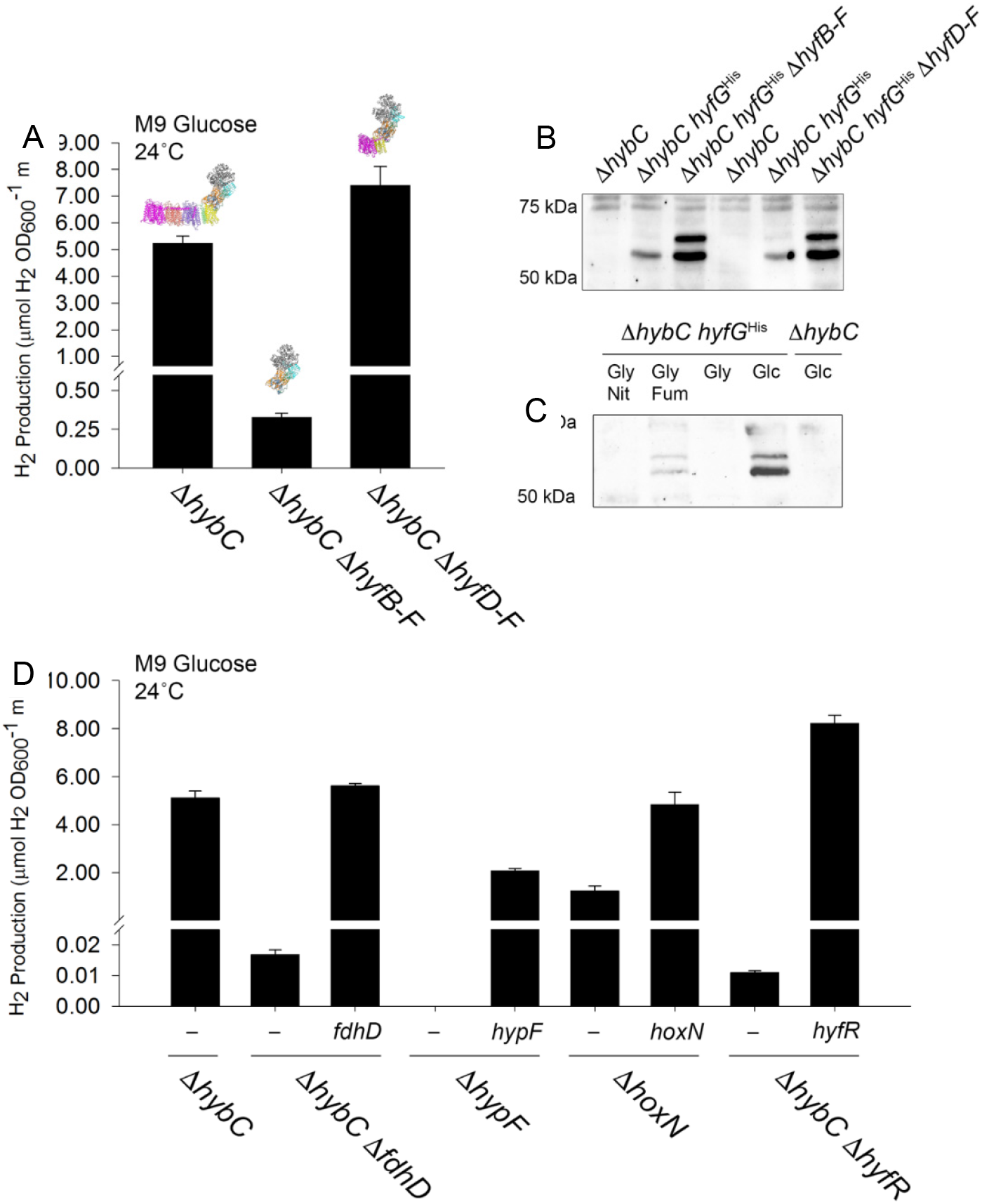
Genetic dissection of FHL-2 activity. **(A)** Some genes encoding the membrane domain are not essential for FHL-2 activity. *P. atrosepticum* strains PH002 (Δ*hybC*), PH009 (Δ*hybC* Δ*hyfB-F*) and PH08 (Δ*hybC* Δ*hyfD-F*) were incubated in M9 medium supplemented with 0.8% (w/v) glucose at 24 °C for 48 hours. **(B)** HyfG^His^ can be detected in strains devoid of membrane subunits. *P. atrosepticum* strains PH002 (Δ*hybC*), PH009 (Δ*hybC*, *hyfG^His^*), PH020 (Δ*hybC hyfG^His^*, Δ*hyfB-F*) and PH021 (Δ*hybC hyfG^His^*, Δ*hyfD-F*) were incubated in M9 medium supplemented with 0.8% (w/v) glucose at 24 °C for 48 hours. Whole cell samples were then prepared by centrifugation, separation of proteins by SDS-PAGE, transfer to nitrocellulose and HyfG^His^ probed with an anti-HIS-HRP antibody. **(C)** HyfG^His^ is induced upon glucose fermentation. Strains PH002 (Δ*hybC*) and PH009 (Δ*hybC*, *hyfG^His^*) were incubated in M9 medium supplemented with either 0.5% (v/v) glycerol and 0.4% (w/v) nitrate (‘Gly Nit’); 0.5% (v/v) glycerol and 0.4% (w/v) fumarate (‘Gly Fum’); 0.5% (v/v) glycerol only (Gly); or 0.8% (w/v) glucose only (‘Glc’) at 24 °C for 48 hours. Whole cell samples were probed for HyfG^His^ with an anti-HIS-HRP antibody. **(D)** The role of accessory genes in FHL-2 activity. Strains PH002 (Δ*hybC*), PH013 (Δ*hybC* Δ*fdhD*), PH010 (Δ*hypF*), PH011 (Δ*hoxN*), and PH015 (Δ*hybC* Δ*hyfR*) were transformed with plasmids encoding either FdhD, HypF, HoxN or HyfR under the control of constitutive promoters. In all cases, the levels of molecular H_2_ in the culture headspace were quantified by GC and normalised to OD_600_ and culture volume. Error bars represent SD (*n* = 3).

Conversely, partial modification of the FHL-2 membrane domain to leave only those subunits present in FHL-1 (Δ*hybC*, Δ*hyfDEF*) had no negative effect on hydrogen production levels (Figure 5A), rather a slight increase was observed. This is consistent with a noticeable enhancement of HyfG^His^ levels in the cells upon removal of the *hyfDEF* genes (Figure 5B). The available evidence suggests that HyfD, HyfE and HyfF have no essential roles in the biosynthesis and hydrogen production activity of FHL-2.

### A requirement for accessory genes in anaerobic hydrogen production

FHL-2 is a multi-subunit metalloenzyme and assembly of such enzymes is often carefully co-ordinated by dedicated chaperones, sometimes called accessory proteins or ‘maturases’. Maturation of molybdenum-dependent formate dehydrogenases has been reported to require the action of an FdhD protein, which is believed to supply an essential sulfur ligand to the active site metal (Arnoux *et al.*, 2015). In *P. atrosecpticum* SCRI1043, *fdhD* (*ECA0093*) is not part of the FHL-2 locus but is located elsewhere on the chromosome next to a gene encoding superoxide dismutase (*sodA* or *ECA0092*) (Bell *et al.*, 2004). Genetic modification of the PH002 strain, containing only Hyd-4 and FHL-2 activity, by the incorporation of a Δ*fdhD* allele (PH013: Δ*hybC*, Δ*fdhD* – Table 1) led to a defect in physiological H_2_ production under fermentative conditions (Figure 5D). This phenotype could be rescued by the provision of extra copies of *fdhD in trans* (Figure 5D).

Maturation of [NiFe]-hydrogenases requires the activity of a network of proteins involved in metal homeostasis and cofactor maturation and insertion (Sargent, 2016). The *P. atrosepticum* FHL-2 locus (Figure 1C) contains a *hoxN* gene (*ECA1252*) encoding a putative membrane-bound nickel ion transporter (Eitinger & Mandrand-Berthelot, 2000). Deletion of the *hoxN* gene in *P. atrosepticum* SCRI1043 (strain PH011, Table 1) reduced hydrogen evolution levels to around 50% of that observed for the parental strain (Figure 5D). Note that there is no other homologue of *hoxN* encoded on the *P. atrosepticum* SCRI1043 genome, but there are several uncharacterised ABC transporters that could be related to the high-affinity *nikA* system (Wu *et al.*, 1991), which could account for the continued availability of nickel for hydrogenase biosynthesis in this experiment.

Once inside the cell, nickel is processed into the Ni-Fe-CO-2CN^−^ cofactor through the action of several enzymes and chaperones. One key step in the biosynthesis of the cofactor is the first step in the generation of CN^−^ from carbamoyl phosphate by HypF (Sargent, 2016). Deletion of the *hypF* gene from *P. atrosepticum* (PH010, Table 1), which is located in the hydrogen metabolism gene cluster under investigation here (Figure 1C), led to the complete abolishment of all detectable H_2_ evolution (Figure 5D). It is the only mutant strain reported here that produces no detectable H_2_ during anaerobic fermentation (Figure 5D). The mutant phenotype could be rescued by supply of *hypF in trans*, but note that full H_2_ evolution levels were not restored (Figure 5D).

Finally, it was observed that a member of the HyfR family of transcriptional regulators was encoded in the hydrogen metabolism gene cluster (Figure 1C). The HyfR protein is predicted to be related to FhlA, which is a formate-sensing transcriptional activator (Skibinski *et al.*, 2002). A Δ*hyfR* strain devoid of the HyfR protein has very low formate hydrogenlyase-2 activity (Figure 5D).

Taken altogether, it can be concluded that all of the genes required for biosynthesis of FHL-2 are functional in *P. atroseptocum* SCRI1043, which is entirely consistent with the physiological data reported here.

## Discussion

### Key differences between FHL-2 and FHL-1

Formate hydrogenlyases can be classified into two structural classes, FHL-1 and FHL-2 (Finney & Sargent, 2019). The most obvious structural difference between an FHL-1, such as the best-characterised *E. coli* enzyme (McDowall *et al.*, 2014, Pinske & Sargent, 2016), and an FHL-2, such as the *P. atrospeticum* enzyme characterised here, is the predicted size and composition of the membrane domain (Figure 1B). The genes encoding FHL-1 include only two membrane proteins, which are a single HycD/HyfC-type protein together with a single HycC/HyfB. This is sufficient to anchor the soluble domain close to the membrane and, in the case of *Thermococcus onnurineus* FHL-1 (Lim *et al.*, 2014) and the related Ech hydrogenase from *Methanosarcina mazei* (Welte *et al.*, 2010), will also allow generation of an ion gradient. Operons encoding FHL-2 complexes encode at least a three further integral membrane proteins. In *P. atrospeticum* these are HyfD and HyfF, which are related to HycC/HyfB, and the HyfE protein. Together, all five proteins are expected to form a stable complex in the membrane, as in the structure of the Group 4 [Ni-Fe]-hydrogenase from *Pyrococcus furiosus* (Yu *et al.*, 2018). Indeed, this large membrane domain is thought to be the ancient progenitor to the ion-pumping membrane domain of respiratory Complex I (Yu *et al.*, 2018, Batista *et al.*, 2013). Given the conservation of these genes, it was surprising that removal of all of the extra membrane proteins from FHL-2 had no discernible effect on the physiological activity of the *P. atrosepticum* system (Figure 5A). In this experiment, the soluble domain was clearly fully assembled and active in formate-dependent hydrogen production (Figure 5A), with Western immunoblotting even pointing towards stabilisation or up-regulation of the catalytic subunit in the absence of *hyfDEF* (Figure 5B). This again highlights the principle of modularity in metalloenzyme evolution, since it is clear that the HyfDEF module may be added or removed depending on both selective pressure and also the, as yet undefined in terms of hydrogenases, biochemical function of these membrane proteins.

Interestingly, removal of the entire membrane domain of FHL-2 (HyfBCDEF) led to a complete loss in formate-dependent hydrogen production, even though the catalytic subunit was synthesised as normal (Figure 5A and B). A similar observation was made with *E. coli* FHL-1, where the catalytic domain was found to be inactive both *in vivo* and *in vitro* when produced in the absence of the membrane domain (Pinske & Sargent, 2016). These data suggest that interaction between the soluble domain and the membrane domain is a key step in the biosynthesis and activation of the enzyme.

The *P. atrosepticum* HyfG catalytic subunit from the Hydrogenase-4 component of FHL-2 shares 74% overall sequence identity (85% similarity) with the *E. coli* HycE protein from Hydrogenase-3/FHL-1. The sequence variation between these two Group 4A hydrogenases is therefore small with only subtle notable differences. For instance, each protein is known or predicted to undergo cleavage during cofactor insertion and maturation leaving a C-terminal arginine residue in the mature form of the proteins. The cleavage sites themselves are slightly differently conserved an FHL-1-type enzyme compared to an FHL-2, for example …R*MTVV… for HycE-like proteins compared to …R*VTLV… for HyfG. This may reflect the need for a different maturation protease for each type of hydrogenase, however this remains to be tested experimentally. In addition, it is notable that both *E. coli* and *P. atrsoepticum hyfG* initiate translation with a GUG start codon.

Phylogenetic analysis of the Group 4A [NiFe]-Hydrogenase subunits, including HycE and HyfG, shows that the enzymes associated with FHL-1 separate into a clearly distinct evolutionary clade from those associated with FHL-2, which form their own distinct clade (**Supp. Figure S2**). Examples of species that encode both FHL-1 and FHL-2 are rare (**Supp. Figure S2**).

### A selenium-free formate dehydrogenase

An in-frame deletion in the *fdhF* gene located in the FHL-2 gene cluster (Figure 1) resulted in a ~500 times reduction in H_2_ production (Figure 4), indicating the majority of H_2_ production from *P. atrosepticum* is dependent on this formate dehydrogenase engaging with Hydrogenase-4 to form an FHL-2 complex. Arguably one of the best-studied FdhF enzymes is the *E. coli* version (Boyington *et al.*, 1997, Gladyshev *et al.*, 1994, Axley *et al.*, 1991). The *E. coli* enzyme contains selenomethionine, which is incorporated co-translationally at a special UGA ‘nonsense’ codon within the coding sequence (Zinoni *et al.*, 1987). Replacement of selenocysteine with cysteine in the *E. coli* enzyme resulted in a dramatically reduced turnover number (Axley *et al.*, 1991). One surprising aspect of *P. atrosepticum* SCRI1043 is that it contains none of the biosynthetic machinery to synthesise selenomethionine (Babujee *et al.*, 2012). The *fdhF* gene studied in this work contains a cysteine codon where selenocysteine would be encoded in the *E. coli* enzyme. Certainly the discovery of a highly active FHL-2 with no need for selenocysteine would benefit scientists interested in engineering this activity into other biological systems.

The FdhF formate dehydrogenase from *P. atrosepticum* shares 65% overall sequence identity (and 85% similarity) with the well-known *E. coli* enzyme (**Supp. Figure S3**). Interestingly, phylogenetic analysis suggests that >50% of species that contain FHL genes utilise a cysteine-dependent, rather than selenocysteine-dependent, formate dehydrogenase (**Supp. Figure S3**). *P. atrosepticum* ECA1507 and ECA1964 were identified here as two FdhF-like proteins that could potentially interact with Hydrogenase-4 to generate novel FHL-like complexes. Sequence analysis revealed ECA1507 and ECA1964 share 65% and 22% overall sequence identity with FdhF, respectively, and phylogenetic analysis determined that ECA1964 is more similar to *E. coli* YdeP than any other predicted molybdenum dependent oxidoreductases in *P. atrosepticum* (**Supp. Figure S3**). YdeP has a putative role in acid resistance in *E. coli*. It is clear that addition of extra copies of ECA1964 in the cell could not complement the Δ*fdhF*, while production of ECA1507 was able to rescue a small amount of FHL-2 activity (Figure 4).

### A role for formate metabolism in a plant pathogen

In the potato pathogen *P. atrosepticum*, FHL-2 activity was found to be expressed at lower growth temperatures (Figure 2). This suggests that FHL-2 may be produced *in planta* during the infection or colonisation event. Formate is produced endogenously by enteric bacteria under fermentative conditions, but plants and tubers have multiple metabolic pathways that generate and consume formate. Potato tubers produce an NAD^+^-dependent formate dehydrogenase (FDH), and the levels of this enzyme are boosted under stress conditions (Hourton-Cabassa *et al.*, 1998). Indeed, proteomic experiments have identified FDH as a differentially-produced protein during wound healing in potato tuber slices, with order of magnitude level changes in protein during this process (Chaves *et al.*, 2009). It could be hypothesised that the expression of FDH in the potato tuber could be coordinated with the initial secretion of formate by a fermenting pathogen. Potentially this would generate NADH from formate in stressed or damaged plant tissues. Recently, it was shown that FDH co-ordinates cell death and defence responses to phytopathogens in *Capsicum annum* (Bell pepper) (Choi *et al.*, 2014). There is also indication that formate and other molecules that lead to the generation of formate, such as methanol and formaldehyde, induce the production of the NAD^+^-dependent FDH, perhaps suggesting there is a signalling response to these C1 compounds in plants (Hourton-Cabassa *et al.*, 1998).

### Concluding remarks

In this work, *P. atrosepticum* SCRI1043 has been established as a tractable new model organism for studying hydrogen metabolism in general and FHL function in particular. The organism is a rare example of a bacterium with an active Hydrogenase-4-containing FHL-2 complex, however, in the course of this work, Hydrogenase-4 activity was reported in *Trabulsiella guamensis*, another γ-Proteobacterium (Lindenstrauss & Pinske, 2019). In *P. atrosepticum*, the active Hydrogenase-4 enzyme operates in tandem with an unusual selenium-free formate dehydrogenase, which may be more amenable to biotechnological engineering than selenium-dependent isoenzymes. In evolutionary terms, the FHL-2 complex has been discussed as a key intermediate in the evolution of the NADH dehydrogenase (Complex I) from a structurally simpler membrane-bound hydrogenase (Schut *et al.*, 2016). The most obvious difference in the predicted quaternary structures inferred from the genetics is the large membrane domain present in FHL-2 compared to FHL-1, and data presented here points to the extra membrane protein being not essential for formate-dependent hydrogen evolution *in vivo*. The role of the FHL membrane domain in generating a transmembrane ion gradient remains to be fully explored in enteric bacteria.

## Methods

### Bacterial strains

The parental *P. atrosepticum* strain used in this study was SCRI1043 (Bell *et al.*, 2004). In-frame deletion and insertion mutants were constructed using pKNG101 suicide vector in *E. coli* strain CC118λ*pir* (Kaniga *et al.*, 1991, Coulthurst *et al.*, 2006). Briefly, upstream and downstream regions (≥600 bp) of the target gene(s) was amplified and inserted into pKNG101 using a three fragment Gibson assembly reaction (HiFi Assembly, NEB). For the insertion of a deca-His encoding sequence into *hyfG*, primers were designed using the NEBuilder online tool to include the deca-His encoding sequence in the overlapping region of the two fragments containing the respective 3’ and 5’ sequences of *hyfG*. After successfully assembly and sequencing of pKNG101 plasmids, the CC118λ*pir* strain with desired plasmid, a HH26 pNJ5000 helper strain, and the desired *P. atrosepticum* strain were grown in rich media, with antibiotics as necessary. Equal volumes of the stationary phase cultures were mixed and 30 µL was spotted on a non-selective rich media plate for 24 hours at 24°C. *P. atrosepticum* cells with the pKNG101 plasmid were initially selected for on minimal media agar with streptomycin (100 µg/ml). After this, single colonies were re-streaked on the fresh minimal media agar with streptomycin. Co-integrants were then grown to stationary phase in rich medium with no selection before the culture was diluted 1/500 with phosphate buffer. Then 30 µL of this diluted culture was plated on minimal media agar with sucrose. These colonies were patch screened for sensitivity to streptomycin before PCR screens were performed to check for presence of the desired mutation(s).

### Plasmids and complementation

All plasmids were cloned using Gibson assembly (HiFi Assembly, NEB) using DNA amplified from *P. atrosepticum* SCRI1043 genomic DNA (Table 1). Genes were cloned into pSU-PROM (Kan^R^), which includes the constitutive *tatA* promoter from *E. coli* (Jack *et al.*, 2004). Complementation plasmids were used to transform electrocompetent *P. atrosepticum* cells using a 2 mm electroporation cuvette (Molecular BioProduct) with application of an electrical pulse (2.5 kV voltage, 25 µF capacitance, 200 Ω resistance and 2mm cuvette length) *via* a Gene Pulsar Xcell electroporator (BioRad). Post recovery, cells were plated on LB Lennox agar plates with 50 µg/ml kanamycin.

### Hydrogen quantification

Hydrogen was directly quantified from 5 mL cultures grown in sealed Hungate tubes (Pinske & Sargent, 2016). Gas-headspace samples were collected using a syringe with Luer lock valve (SGE), Samples were analysed using Gas Chromatography (Shimadzu GC-2014, capillary column, TCD detector). A hydrogen standard curve was used to quantify sample hydrogen content, this was then normalised to optical density (OD_600_) and culture volume (Pinske & Sargent, 2016).

### Western immunoblotting

Proteins samples were first separated by SDS-PAGE using the method of Laemmli (Laemmli, 1970) before transfer to nitrocellulose (Dunn, 1986). Nitrocellulose membranes were challenged with an anti-His-HRP antibody (Alpha Diagnostics) and a GeneGnome instrument (SynGene) was used to visualise immunoreactive bands following addition of ECL reagent (Bio-Rad).

### Structure modelling and phylogenetic analysis

Structural modelling of the formate hydrogenlyases complexes was performed using Phyre^2^ predictions of respective subunits (Kelley & Sternberg, 2009). Using Chimera (Pettersen *et al.*, 2004) the X-ray crystal structure of *Thermus thermophilus* Respiratory Complex I (4HEA) and the Cryo-EM structure of Membrane Bound Hydrogenase (6CFW), the individual FHL-2 subunits were manually assembled into a putative complex organisation for FHL-1 and FHL-2. Phylogenetic analysis of *E. coli* FdhF-like proteins from organisms possessing a Group 4A [NiFe]-hydrogenase utilised the HydDB database (Greening *et al.*, 2015) to collect accession numbers for all [NiFe]-hydrogenase subunits. In each organism the FdhF orthologs were identified before MUSCLE multiple sequence alignment in Jalview (Waterhouse *et al.*, 2009). Through percentage identity tree generation and manual inspection the closest FdhF-like proteins in each organism were identified. FigTree (http://tree.bio.ed.ac.uk/software/figtree) was used to visualise the finalised phylogenetic trees.

## Acknowledgements

This research was funded primarily by the BBSRC through award of an EASTBIO PhD studentship to AJF (1510231). MA was funded by the European Union Horizon 2020 Research & Innovation Programme under the Marie Skłodowska-Curie Grant agreement 654006. SJC is a Wellcome Trust Senior Research Fellow.

