## Supplementary Figures for "The plant pathogen *Pectobacterium atrosepticum* contains a functional formate hydrogenlyase-2 complex"

**SUPPLEMENTARY INFORMATION**

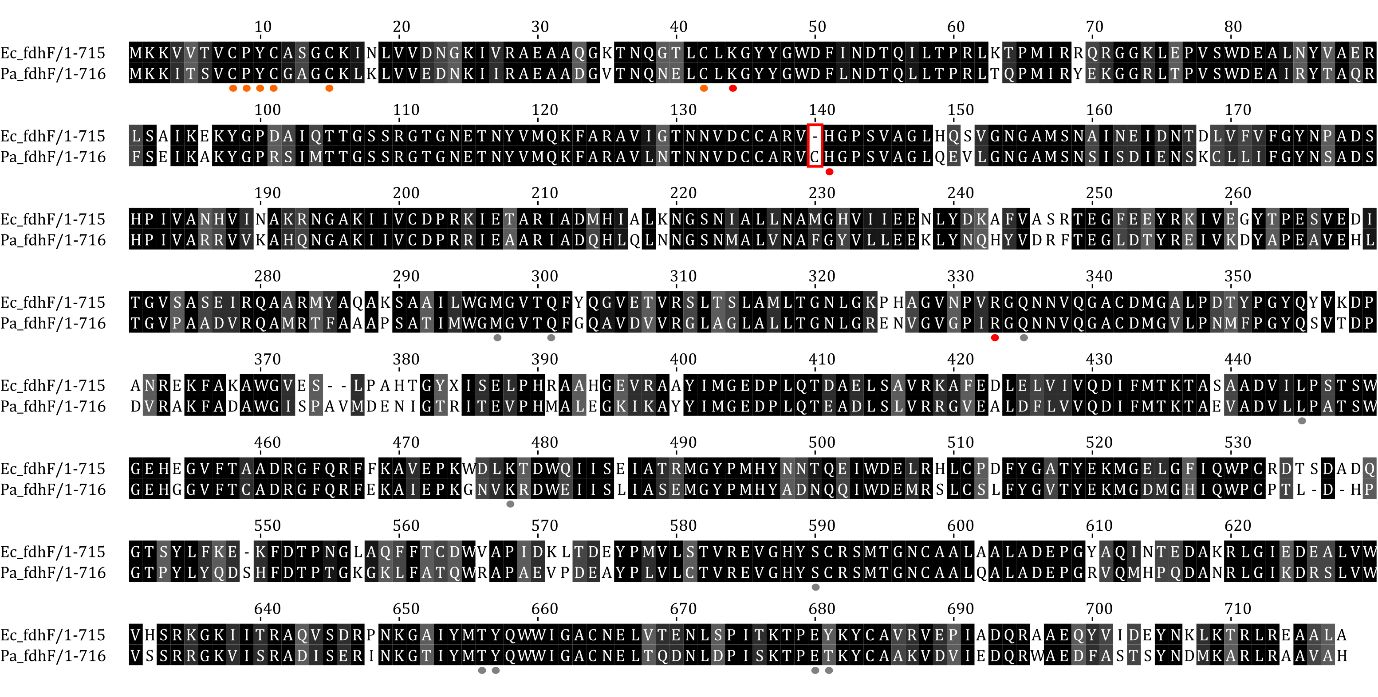

**Supp Figure S1: A selenocysteine-free formate dehydrogenase in *P. atrosepticum***

Primary sequence alignment of *E. coli* FdhF and *P. atrosepticum* SCRI1043 FdhF (ECA1250) using Jalview. Orange annotations show Fe-S cluster co-ordination residues, Red annotations show active site residues and Grey annotations show Mo-*bis*MGD co-ordination – all based on the *E. coli* FdhF crystal structure (1AA6). A red box highlights the selenocysteine/cysteine key difference between the two homologs.

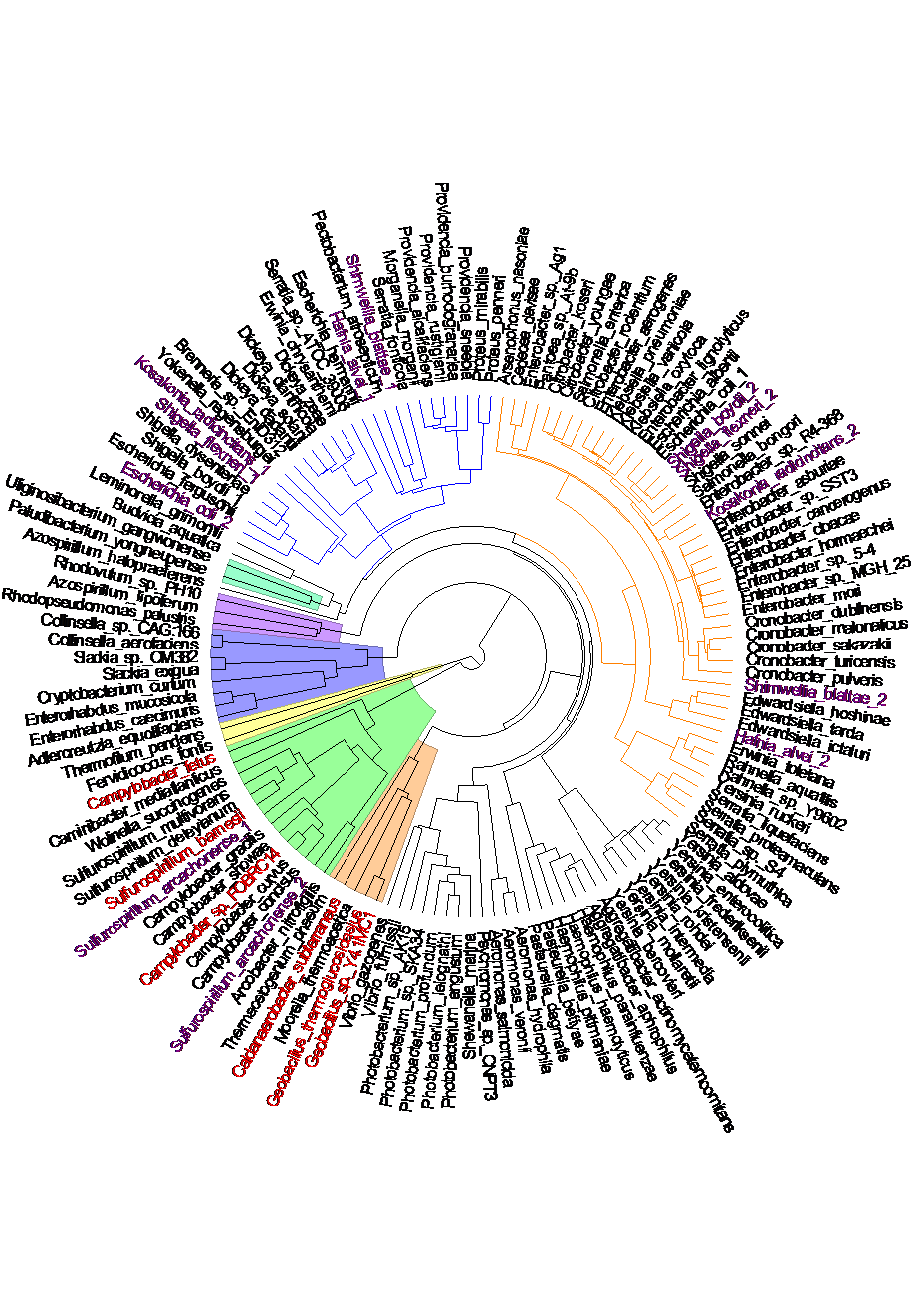

**Supp Figure S2: Phylogenetic analysis of Group 4A [NiFe] hydrogenase catalytic subunits**

A phylogenetic tree of Group4A [NiFe]-hydrogenase catalytic subunits. Proteins linked by the orange line are those predicted to be of the HycE-type (Hyd-3), while those linked by the blue line are those predicted to be of the HyfG type (Hyd-4). The coloured sections represent different phylogenetic groups: brown section are Firmicutes (five members represented); green section are the Epsilonproteobacteria (14); yellow section are the Crenarchaeota (Archaea) (two); lilac section are Actinobacteria (eight); purple section are Alphaproteobacteria (three + *Azospirillum halopraeferens*); light blue section are Betaproteobacteria (two); and those with no highlight colour are the Gammaproteobacteria. Species names highlighted in purple text have genomes where no FdhF (formate dehydrogenase) more than one Group4A hydrogenase was identified, and species names highlighted in red text have genomes where no FdhF (formate dehydrogenase) was identified.

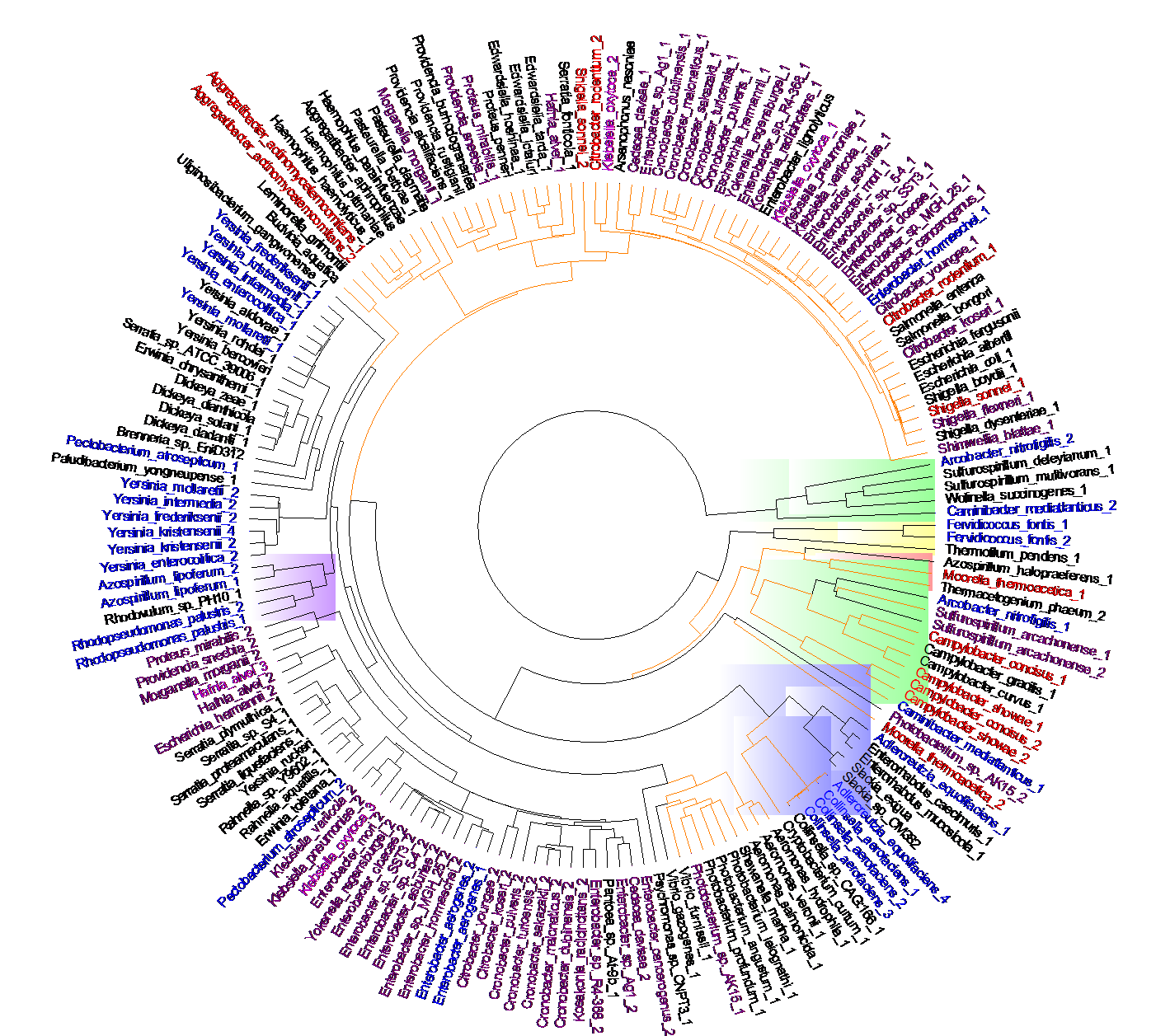

**Supp Figure S3: Phylogenetic analysis of metal-dependent formate dehydrogenases in prokaryotes also containing a Group 4A [NiFe]-hydrogenase catalytic subunit.**

A phylogenetic tree of FdhF-like formate dehydrogenase catalytic subunits. Proteins linked by the orange line are those predicted to contain an active site seleno-cysteine residue, while those linked by the black line are those predicted to contain an active site cysteine. The coloured sections represent different phylogenetic groups: brown section are Firmicutes; green section are the Epsilonproteobacteria; yellow section are the Crenarchaeota (Archaea); lilac section are Actinobacteria; purple section are Alphaproteobacteria; and those with no highlight colour are the Gammaproteobacteria. Species names highlighted in red text have genomes where two FdhF (seleno-Cys) can be identified; species names highlighted in purple text have genomes where two FdhF (one seleno-Cys, one Cys) can be identified; species names highlighted in pink text have genomes where more than two FdhF (mixture of seleno-Cys and Cys) can be identified; and species names highlighted in blue text have genomes where two FdhF (both Cys) can be identified.

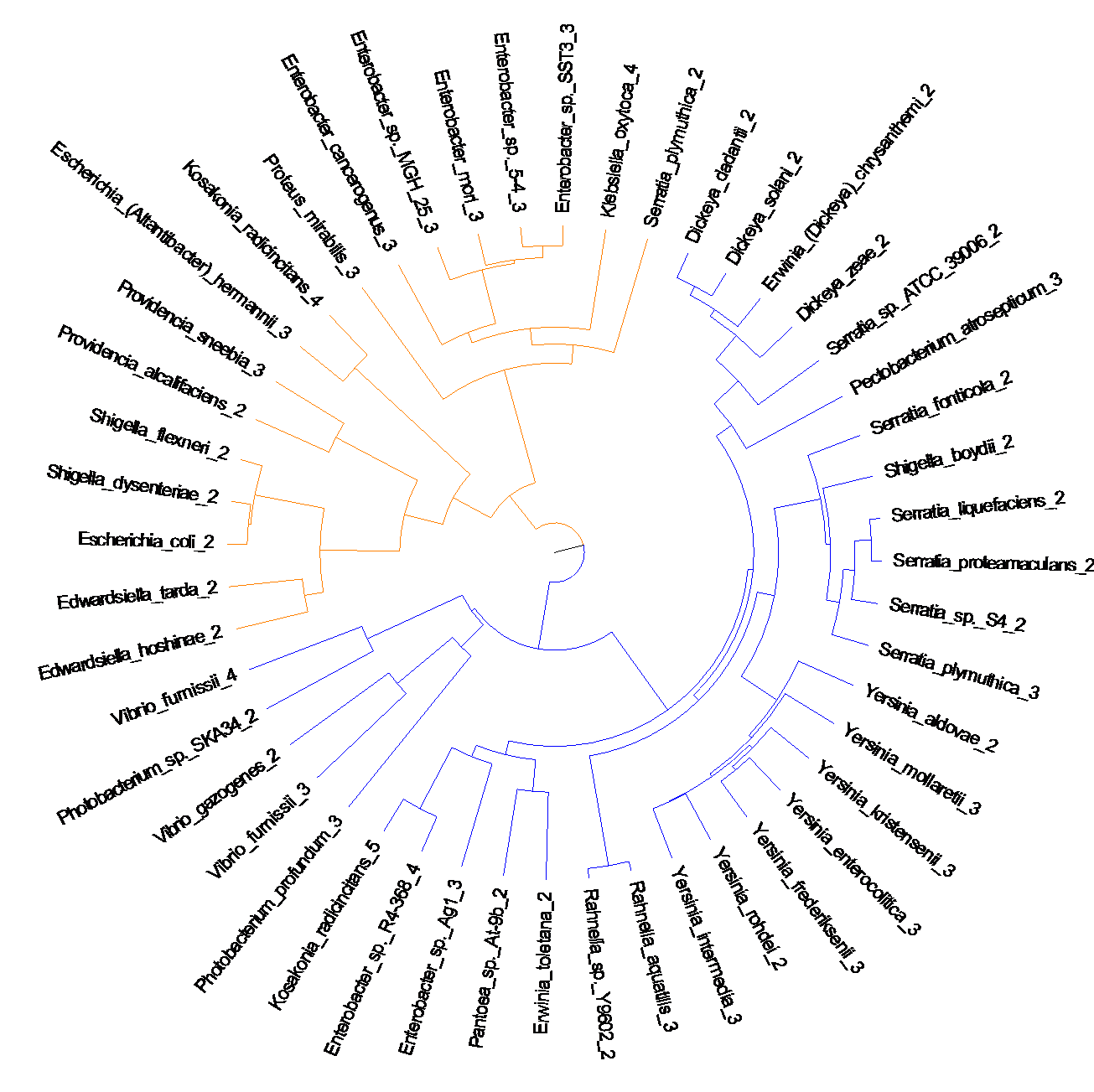

**Supp Figure S4: Phylogenetic analysis of YdeP proteins in prokaryotes.**

A phylogenetic tree of YdeP-like Mo/W-dependent catalytic subunits, which are related to formate dehydrogenases. Two separate clades are evident.
